# Evolution and cell-type specificity of human-specific genes preferentially expressed in progenitors of fetal neocortex

**DOI:** 10.1101/225722

**Authors:** Marta Florio, Michael Heide, Holger Brandl, Anneline Pinson, Sylke Winkler, Pauline Wimberger, Wieland B. Huttner, Michael Hiller

## Abstract

To understand the molecular basis underlying the expansion of the neocortex during primate, and notably human, evolution, it is essential to identify the genes that are particularly active in the neural stem and progenitor cells of developing neocortex. Here, we have used existing transcriptome datasets to carry out a comprehensive screen for protein-coding genes preferentially expressed in progenitors of fetal human neocortex. In addition to the previously studied gene *ARHGAP11B*, we show that ten known and two newly identified human-specific genes exhibit such expression, however with distinct neural progenitor cell-type specificity compared to their ancestral paralogs. Furthermore, we identify 41 additional human genes with progenitor-enriched expression which have orthologs only in primates. Our study not only provides a resource of genes that are candidates to exert specific, and novel, roles in neocortical development, but also reveals that distinct mechanisms gave rise to these genes during primate, and notably human, evolution.

## Introduction

The expansion of the neocortex in the course of human evolution provides an essential basis for our cognitive abilities (Azevedo et al., 2009; Borrell and Reillo, 2012; Buckner and Krienen, 2013; Dehay et al., 2015; Florio and Huttner, 2014; Kaas, 2013; Lui et al., 2011; Namba and Huttner, 2017; Rakic, 2009; Sousa et al., 2017; Striedter, 2005). This expansion ultimately reflects an increase in the proliferative capacity of the neural stem and progenitor cells in the developing human neocortex (from now on collectively referred to as cortical neural progenitor cells, cNPCs) (Azevedo et al., 2009; Bae et al., 2015; Borrell and Reillo, 2012; Dehay et al., 2015; Florio and Huttner, 2014; Lui et al., 2011; Namba and Huttner, 2017; Rakic, 2009), as well as in the duration of their proliferative, neurogenic and gliogenic phases (Lewitus et al., 2014; Otani et al., 2016). It is therefore a fundamental task to elucidate the underlying molecular basis, that is, the changes in our genome that endow human cNPCs with these neocortical expansion-promoting properties.

One approach towards this goal is to identify which of the genes that are particularly active in human cNPCs exhibit a human-specific expression pattern, or even are human-specific. We previously isolated, and determined the transcriptomes of, two major cNPC types from embryonic mouse and fetal human neocortex (Florio et al., 2015), (i) the apical (or ventricular) radial glia (aRG), the primary neuroepithelial cell-derived apical progenitor type (Götz and Huttner, 2005; Kriegstein and Götz, 2003), and (ii) the basal (or outer) radial glia (bRG), the key type of basal progenitor implicated in neocortical expansion (Betizeau et al., 2013; Borrell and Götz, 2014; Borrell and Reillo, 2012; Florio and Huttner, 2014; Lui et al., 2011). This led to the identification of 263 protein-coding human genes that are much more highly expressed in human bRG and aRG than in a neuron-enriched fraction (Florio et al., 2015). Of these, 207 genes have orthologs in the mouse genome but are not expressed in mouse cNPCs, whereas 56 genes lack mouse orthologs. Among the latter, the gene with the highest specificity of expression in bRG and aRG was found to be *ARHGAP11B*, a human-specific gene (Antonacci et al., 2014; Dennis et al., 2017; Riley et al., 2002; Sudmant et al., 2010) that we showed to be capable of basal progenitor amplification in embryonic mouse neocortex and to likely have contributed to the evolutionary expansion of the human neocortex (Florio et al., 2015; Florio et al., 2016).

Our previous finding that, in addition to *ARHGAP11B*, there are 55 other human genes without mouse orthologs that are predominantly expressed in bRG and aRG (Florio et al., 2015), raises the possibility that some of these genes may be human-specific and may affect the behaviour of human cNPCs. To investigate the evolution and cell-type specificity of expression of such genes, we have now data-mined our previous dataset (Florio et al., 2015) as well as three additional ones (Fietz et al., 2012; Miller et al., 2014; Pollen et al., 2015) to carry out a comprehensive screen for protein-coding genes preferentially expressed in cNPCs of fetal human neocortex. We find that, in addition to *ARHGAP11B*, 12 other human-specific genes (10 previously and 2 newly identified ones) show preferential expression in cNPCs. Furthermore, we identify 41 additional human genes exhibiting such expression for which orthologs are found in primate but not in non-primate mammalian genomes. We provide information on the evolutionary mechanisms leading to the origin of several of these primate-specific genes, including gene duplication and transposition. Moreover, we analyze the cell-type specific expression of most of the human-specific genes, including their splice variant expression patterns. Finally, by comparing the expression of the human-specific genes with their respective ancestral paralog, we show a substantial degree of gene expression divergence upon gene duplication, suggesting potential neofunctionalization. Our study thus provides a resource of genes that are candidates to exert specific roles in the development and evolution of the primate, and notably human, neocortex.

## Results

### Screen of distinct transcriptome datasets from fetal human neocortex for protein-coding genes preferentially expressed in neural stem and progenitor cells

To identify genes preferentially expressed in the cNPCs of the fetal human neocortex, we analyzed four distinct, published transcriptome datasets obtained from human neocortical tissue ranging from 13 to 19 weeks post conception (wpc). First, the genome-wide RNA-Seq dataset obtained from specific neocortical zones isolated by laser capture microdissection (LCM) (Fietz et al., 2012), which we screened for all protein-coding genes that are more highly expressed in the VZ, iSVZ and/or oSVZ than the cortical plate (CP) (as determined by differential gene expression (DGE) analysis, p <0.01). This yielded 2758 genes (Fig. 1A, B). Second, the Allen Brain Institute microarray dataset obtained from LCM-isolated specific neocortical zones (Miller et al., 2014), which we screened for all protein-coding genes with positive laminar correlation (correlation coefficient >0.5) with either the VZ, iSVZ or oSVZ as compared to the zones enriched in postmitotic cells (intermediate zone (IZ), subplate, CP, marginal zone, subpial granular zone). This yielded 4555 genes (Fig. 1A, B). Third, the genome-wide RNA-Seq dataset obtained from specific neocortical cell types isolated by fluorescence-activated cell sorting (FACS) (Florio et al., 2015), which we screened for all protein-coding genes more highly expressed (as determined by DGE analysis, p<0.01) in aRG and/or bRG in S-G2-M as compared to the cell population enriched in postmitotic neurons but also containing bRG in G1. This yielded 2106 genes (Fig. 1A, B). Fourth, the dataset obtained from genome-wide single-cell RNA-Seq of dissociated cells captured from microdissected VZ and SVZ (Pollen et al., 2015), which we screened for all protein-coding genes positively correlated with either radial glial cells, bIPs or both (correlation coefficient >0.1) and negatively correlated with neurons (correlation coefficient <0.1). This yielded 5335 genes (Fig. 1A, B).

**Fig. 1.**
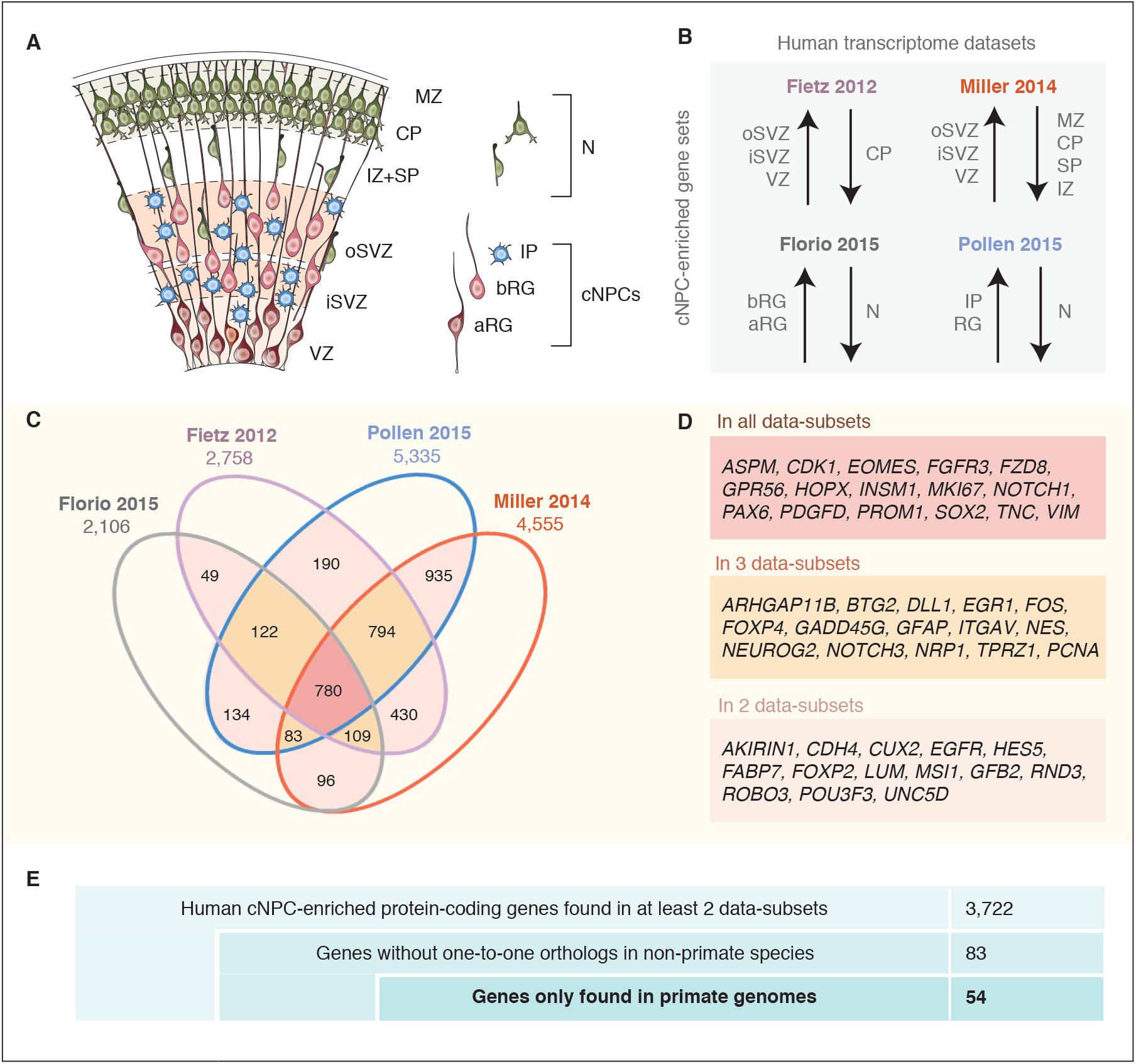
A screen for human cNPC-enriched protein-coding genes and determination which of them have orthologs only in primates. (**A**) Cartoon illustrating the main zones and neural cell types in the fetal human cortical wall that were screened for differential gene expression in the human transcriptome datasets as depicted in (B). Adapted from (Florio et al., 2017). SP, subplate; MZ, marginal zone. (**B**) The indicated four published transcriptome datasets from fetal human neocortical tissue (Fietz et al., 2012; Miller et al., 2014) and cell populations (Florio et al., 2015; Pollen et al., 2015) were screened for protein-coding genes showing higher levels of mRNA expression in the indicated germinal zones and cNPC types than in than in the non-proliferative zones and neurons. (**C**) Venn diagram showing the data-subsets of human protein-coding genes displaying the differential gene expression pattern depicted in (B). Numbers within the diagram indicate genes found in two (pink), three (yellow) or all four (red) data-subsets. Genes found in at least two data-subsets were considered as being cNPC-enriched. (**D**) Selected genes with established biological roles found in two (pink), three (yellow), or all four (red) data-subsets. (**E**) Stepwise analysis leading from the 3,722 human cNPC-enriched protein-coding genes to the identification of 54 primate-specific genes.

Next, we determined how many of the protein-coding genes exhibiting the above described differential expression pattern were found in all four datasets. This was the case for 780 genes (Fig. 1C, red). We also determined the number of genes found in three of the four datasets (four combinations, Fig. 1C orange), and of those found in two datasets (six combinations, Fig. 1C pink). Together this yielded a catalogue of 3,722 human genes with preferential expression in cNPCs (from here on referred to as cNPC-enriched genes) (see Table S1).

These 3,722 genes included well-known molecular players involved in cNPC function and established markers of cNPCs, notably radial glia, (e.g. *EOMES, FABP7, GFAP, HOPX, NES, PAX6, SOX2, VIM*), cell proliferation (e.g. *MKI67, PCNA*), Notch signaling (*DLL1, HES5*), and extracellular matrix and growth factor signaling (e.g. *FGFR3, ITGAV, LUM, TNC*) (listed in Fig. 1D). Moreover, several genes recently implicated in human-specific aspects of cNPC proliferation and neocortex formation (Florio et al., 2017; Mitchell and Silver, 2017; Sousa et al., 2017) (e.g. *ARHGAP11B, FOXP2, FZD8, GPR56, PDGFD*) were found in the analyzed datasets, though not necessarily in all four (Fig. 1D). The latter finding, on the one hand, likely reflects the diversity of the cNPC enrichment strategies and mode of transcriptome analysis adopted to obtain the four datasets, and on the other hand highlights the significance of data-mining all these datasets in combination. On a general note, the catalog of 3,722 cNPC-enriched human genes presented here (Table S1) provides an integrative and methodologically unbiased tool to interrogate the cNPC enrichment of candidate genes of interest, and to potentially uncover new genes involved in cNPC function during fetal human corticogenesis.

### Identification of primate-specific genes

Primate-specific, notably human-specific, genes expressed in cNPCs have gained increasing attention for their potential role in species-specific aspects of neocortical development, including neurogenesis (Charrier et al., 2012; Dennis et al., 2017; Florio et al., 2017; Sousa et al., 2017). To determine how many of the 3,722 human cNPC-enriched protein-coding genes had orthologs only in primates but not in non-primate species, we eliminated from this gene set all those genes with an annotated one-to-one ortholog in any of the sequenced non-primate genomes (Fig. 1E). This greatly reduced the number of genes from 3,722 to 83 genes.

Next, we examined these 83 genes to extract those that are truly primate-specific. By inspecting genomic alignments, gene neighborhoods and gene annotations in primate and non-primate mammals, we concluded that 29 of these genes likely have an ortholog in non-primate mammals and we therefore excluded them from further analysis. The remaining 54 genes were considered to be truly primate-specific (Fig. 1E) and are of special evolutionary interest because of their orthology to the human cNPC-enriched protein-coding genes.

### Phylogenetic analysis of the primate-specific genes

To trace the phylogeny of the 54 primate-specific genes and to infer their ancestry, we investigated in which species these genes exhibit an intact reading frame, and used this information to assign each gene to a primate clade. First, we found that 28 of these 54 genes exist in the genomes of tarsiers, monkeys and apes (though not necessarily in every species of these clades) and thus likely predate the ape (Hominoidea) ancestor that lived ~28 Mya (Kumar et al., 2017) (Fig. 2). Specifically, some of these 28 genes can already be detected in haplorrhini, some in simiiformes and some at first in catarrhini (Fig. 2). Strikingly, more than half of these 28 genes encode zinc finger proteins (Table 1). Second, we found that of the remaining 26 primate-specific genes that exist only in the genomes of apes (hominoidea), although not necessarily in every ape species (Fig. 2), 13 genes exist not only in the human genome but also in non-human ape genomes. It is interesting to note that 11 of these 13 genes exist only in great apes (hominidae) and thus likely arose after the lineage split (~18 Mya, Kumar et al., 2017) leading to the gibbon vs. the great apes, and 8 of these 11 genes do not exist in the orangutan and thus likely arose after the great ape ancestor split into the orangutan and Homininae lineage ~15 Mya (Kumar et al., 2017) (Fig. 2). Of the two hominoidea-specific genes already detected in the gibbon, *PTTG2* deserves special comment and is further discussed below.

**Fig. 2.**
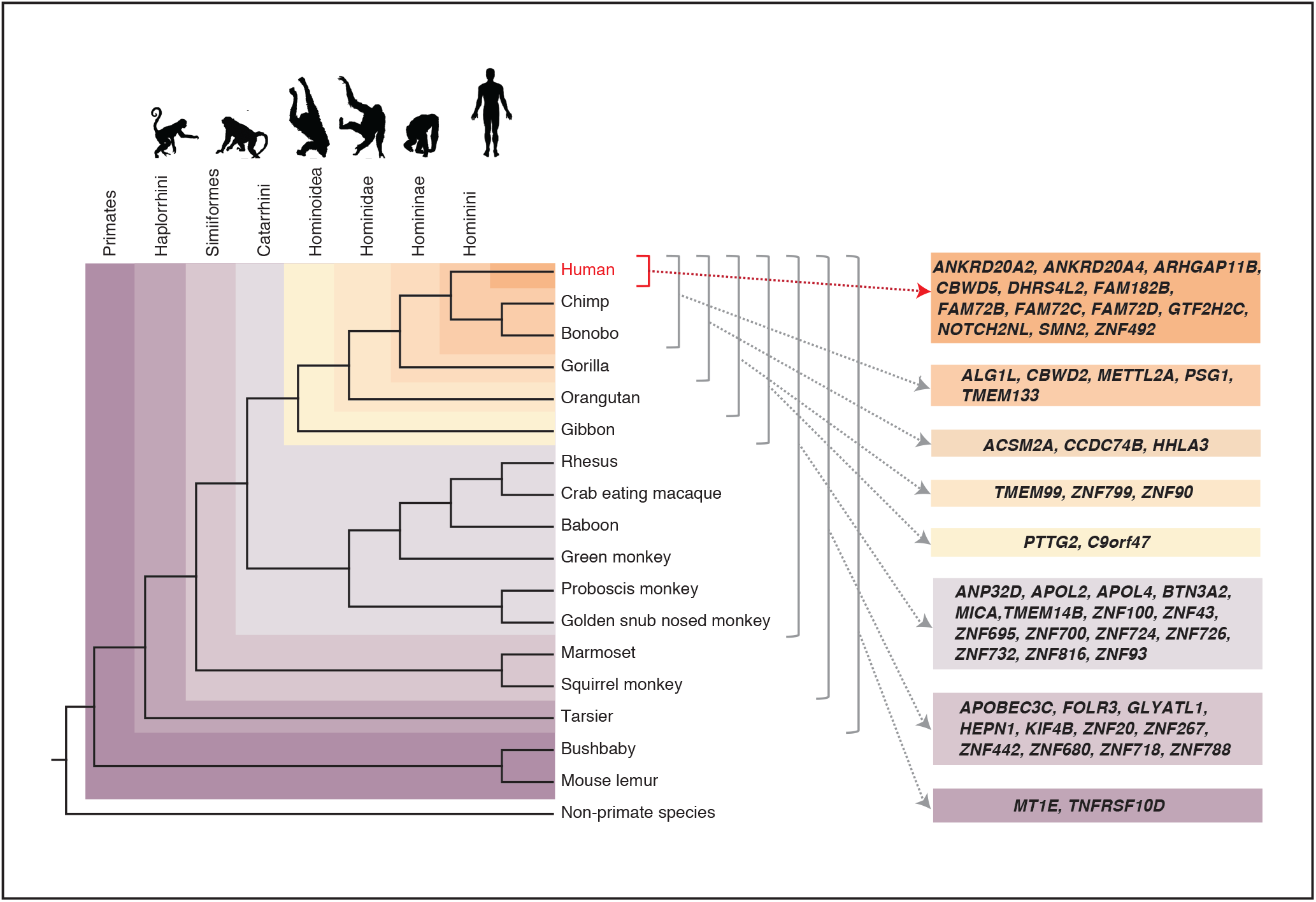
Occurrence of the primate-specific genes in the various primate clades. Assignment of the 54 primate-specific genes to a primate clade, based on the primate genome(s) in which an intact reading frame was found in the present analysis. Clades are specified on the top left. The color-coding and brackets indicate the species in each clade analyzed in the present study. Note that the occurrence of the genes in the various clades does not necessarily apply to every species in the clade.

Finally, we found that 13 of the 54 primate-specific genes were only present in the human genome, and thus arose (or evolved to their present state) in the human lineage after its split from the lineage leading to the chimpanzee (Fig. 2) (~5-7 Mya, Brunet et al., 2005; Brunet et al., 2002; Vignaud et al., 2002). These 13 human-specific genes include *ARHGAP11B*, a gene that we reported previously to have a key role in cNPC proliferation and neocortex expansion (Florio et al., 2015; Florio et al., 2016) and that was also present in the archaic genomes of Neandertals and Denisovans (Antonacci et al., 2014; Florio et al., 2015; Meyer et al., 2012; Prüfer et al., 2014; Sudmant et al., 2010). Similar to *ARHGAP11B*, 11 of the remaining 12 human-specific genes existed also in the genomes of Neandertals and Denisovans (Dennis et al., 2017; Sudmant et al., 2010) and present data, see Table 1) and thus arose before the split of the lineages leading to modern humans vs. Neandertals/Denisovans ~500,000 years ago (Meyer et al., 2012; Prüfer et al., 2014). Of note, *SMN2* is the only gene in this set that has been reported to have arisen in the lineage leading to modern humans after its divergence from the lineage leading to Neandertals and Denisovans (Dennis et al., 2017).

### Analysis of the evolution of selected primate-specific genes reveals distinct mechanisms

Next, we sought to determine how these 54 primate-specific genes evolved. With regard to the primate-specific genes that are not human-specific, we focused on three genes that we selected in light of their potential biological role, *MICA, KIF4B* and *PTTG2*.

*MICA* (*MHC class I polypeptide-related sequence A*) is a paradigmatic example of a gene arising by gene duplication (Bailey et al., 2002; Eichler et al., 2004; Fortna et al., 2004; Hurles, 2004), a well-known driving force of genome evolution (Lynch and Conery, 2000). *MICA* arose by duplication of the widely occurring *MICB* gene. As *MICA* is found in the genomes of apes and Old-World monkeys (Catarrhini), but not New-World monkeys (Fig. 2), this gene duplication presumably occurred after the separation of the lineages leading to New-World monkeys vs. and Catarrhini ~47 Mya (Kumar et al., 2017). *MICA* is the only gene among the 54 primate-specific genes analyzed in the present study that has an established relationship to the MHC locus (Bahram et al., 1994), pointing to a possible primate-specific interaction between cNPCs and cells of the immune system.

Besides gene duplication, however, other mechanisms were found to underlie the evolution of primate-specific genes. A notable example is *KIF4B* (*Kinesin Family Member 4B*), a gene encoding a kinesin involved in spindle organization during cytokinesis (Zhu et al., 2005). In fact, *KIF4B* is the only member of the kinesin superfamily among the 54 primate-specific genes. *KIF4B* is specific to apes, Old-World monkeys and New-World monkeys (Simiiformes, Fig. 2) and evolved by retroposition of *KIF4A*, a gene with a near-ubiquitous occurrence in the animal kingdom (Hirokawa et al., 2009). This retroposition involved the reverse transcription of a spliced *KIF4A* mRNA followed by insertion of the DNA into the genome as an intronless copy of *KIF4A*.

Similar to *KIF4B*, the primate-specific gene *PTTG2 (pituitary tumor transforming 2*) arose by retroposition. Specifically, *PTTG2* arose by reverse transcription of the spliced mRNA of *PTTG1*, a gene encompassing five protein-coding exons conserved in reptiles, birds and mammals and implicated in promoting proliferation of pituitary tumor cells (Dominguez et al., 1998; Vlotides et al., 2007; Zhang et al., 1999). However, while *KIF4B* inserted into an intergenic locus (that however allowed its transcription), the intron-less protein-coding *PTTG2* inserted into intron 2 of the *TBC1D1* gene (Fig. 3A), which encodes a Rab-GTPase activating protein (Roach et al., 2007). This *PTTG2* retroposition event presumably occurred in the ancestor of New-World monkeys, Old-World monkeys and apes (simiiformes). Remarkably, after retroposition, the *PTTG2* gene underwent two principally different lines of evolution. In all non-hominoidea simiiformes (see Fig. 2), consistent with neutral evolution, *PTTG2* accumulated frameshifting deletions and translational stop codon mutations that cause premature termination of the open reading frame (Fig. 3B). In contrast, in hominoidea (apes and humans, see Fig. 2), the *PTTG2* reading frame remained open, with one noticeable change. This is a 1-bp insertion (T, see Fig. 3A) near the 3’ end of the *PTTG2* open reading frame that causes a shift in the reading frame, resulting in a new 13-amino acid-long C-terminal sequence of PTTG2 in great apes (including human) (as opposed to 24 amino acids in PTTG1) (Fig. 3B). This PTTG2-specific sequence lacks the cluster of acidic residues found in the C-terminal sequence of PTTG1. In the case of the gibbon, however, the *PTTG2* gene carries (in addition to the 1-bp T insertion) a 22-bp deletion a few nucleotides 5’ to this insertion. This causes yet another shift in the reading frame that results in the replacement of the C-terminal 25 amino acids of the PTTG2 of great apes (including human) by an 18-amino acid-long sequence (Fig. 3B). The potential consequences of these changes in protein sequence for the function of PTTG2 with regard to cell proliferation are discussed below.

**Fig. 3.**
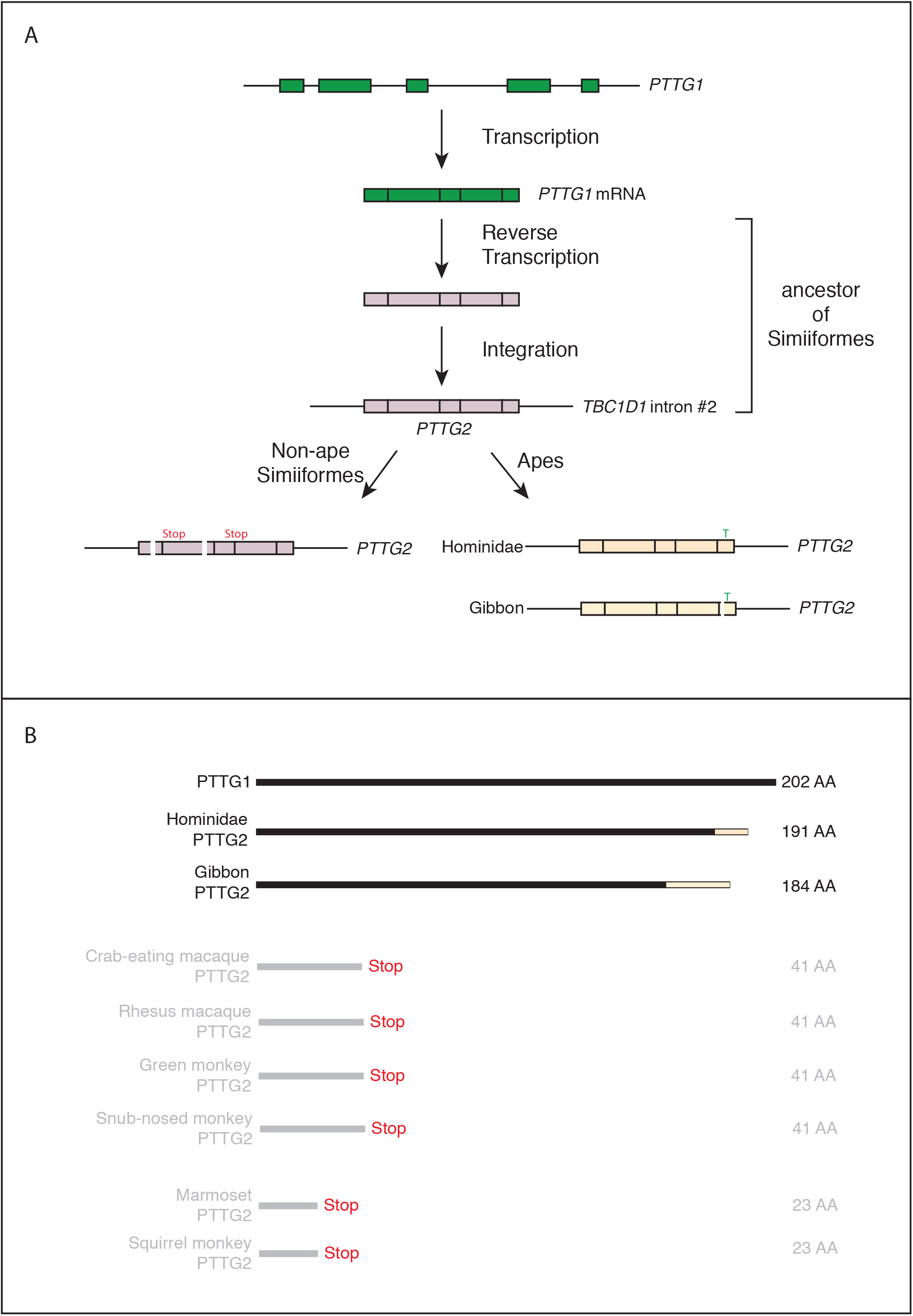
Evolutionary origin of the *PTTG2* gene. (**A**) Origin of the *PTTG2* gene by reverse transcription of the *PTTG1* mRNA and insertion as a retroposon into the *TBC1D1* locus in the ancestor to New-World monkeys, Old-World monkeys and apes (Simiiformes). (**B**) Comparison of the PTTG1 and hominoidea PTTG2 polypeptides, and of the prematurely closed open reading frames of non-ape simiiformes *PTTG2*.

### A variety of evolutionary mechanisms gave rise to the human-specific cNPC-enriched protein-coding genes

We next investigated how the 13 human-specific cNPC-enriched protein-coding genes evolved. Nine of them arose by duplications of entire genes (Bailey et al., 2002; Eichler et al., 2004; Fortna et al., 2004; Hurles, 2004) (Figure 4A). Two arose by partial gene duplication. These genes are *ARHGAP11B* and *NOTCH2NL* (Antonacci et al., 2014; Dennis et al., 2017; Dougherty et al., 2017; Riley et al., 2002). *ARHGAP11B* arose from partial duplication of *ARHGAP11A*, which encodes a Rho GTPase activating protein (RhoGAP). *ARHGAP11B* comprises the entire GAP domain of ARHGAP11A but, due to a single-base substitution (C–>G) creating a new splice donor site, encodes a protein with a truncated GAP domain followed by a unique, human-specific C-terminal amino acid sequence (Florio et al., 2015; Florio et al., 2016) (Fig. 4B).

**Fig. 4.**
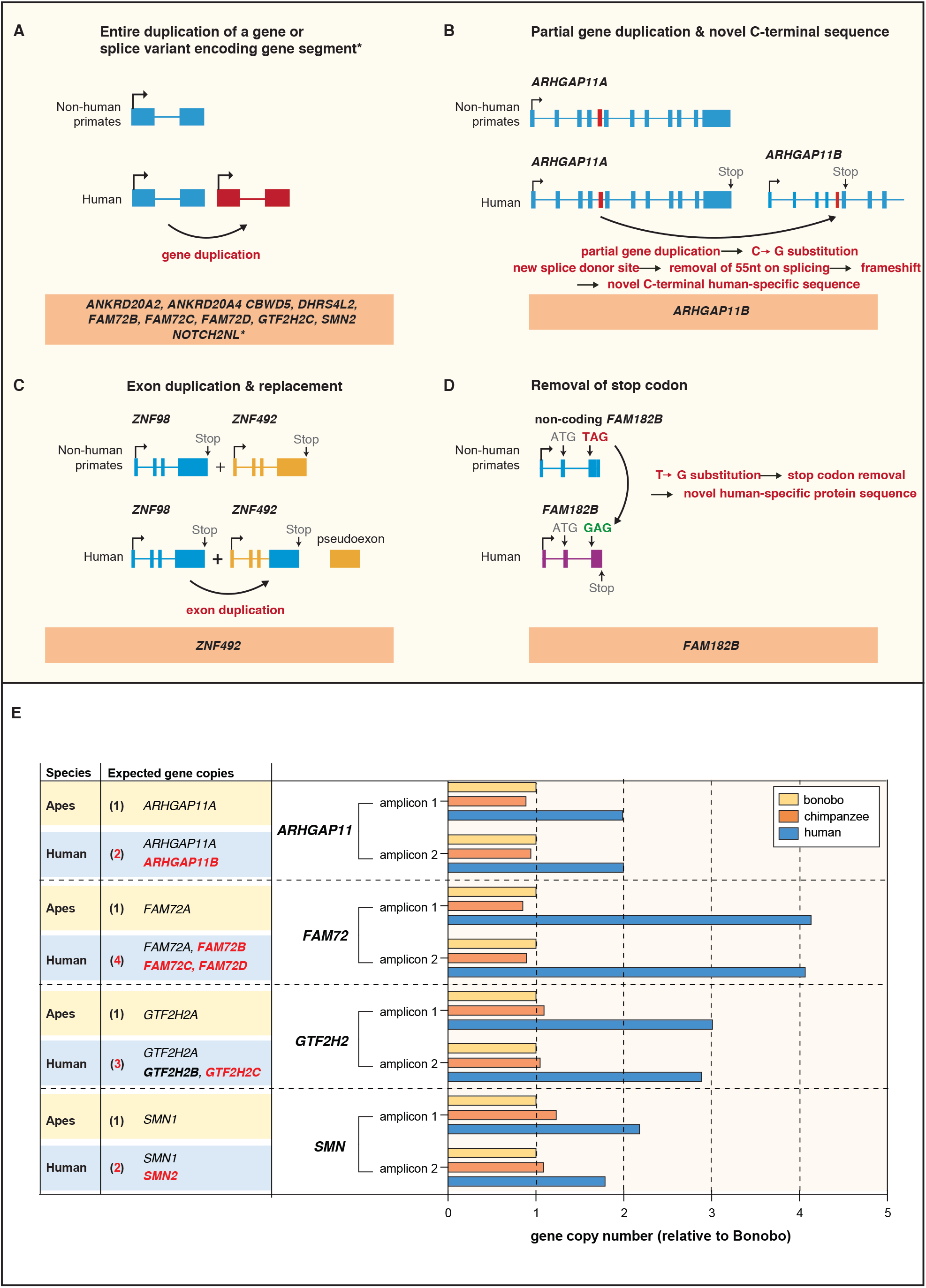
Evolution of the human-specific cNPC-enriched protein-coding genes. Diagrams depicting the evolutionary origin of the 13 human-specific genes. (**A**) Duplication of the entire ancestral gene, which applies to nine of the human-specific genes. *NOTCH2NL* is included in this group because it arose by entire duplication of a short *NOTCH2* splice variant; the latter, however, comprises only parts of the *NOTCH2* gene (hence the asterisk). Note that the gene duplication giving rise to *SMN2* occurred after the Neandertal – modern human lineage split, whereas the other eight gene duplications occurred before that split (Dennis et al., 2017). (**B**) Partial gene duplication (~5 Mya) giving rise to *ARHGAP11B* (Antonacci et al., 2014; Dennis et al., 2017; Riley et al., 2002). Note that a single C–>G substitution in exon 5 (red box), which likely occurred after the gene duplication event but before the Neandertal – modern human lineage split, created a new splice donor site, causing a reading frame shift that resulted in a novel, human-specific 47 amino acid C-terminal sequence (Florio et al., 2015; Florio et al., 2016). (**C**) Exon duplication and replacement giving rise to human *ZNF492*. Exon 4 of *ZNF98* (blue) is duplicated and inserted into intron 3 of *ZNF492* (orange), rendering the original *ZNF492* exon 4 a pseudoexon. (**D**) Removal of a stop codon converting the non-coding *FAM182B* of non-human primates into the protein-coding human *FAM182B*. A single T–>G substitution removes the stop codon at the 5' end of exon 3, thereby generating an open reading frame (purple). (**E**) Validation of the human-specific nature of selected human genes by determination of their copy numbers. Human (blue), chimpanzee (orange) and bonobo (yellow) genomic DNA was used as template to perform a qPCR that would generate two distinct amplicons of both, the gene common to all three species (black regular letters) and the human-specific gene(s) under study (red bold letters), as indicated. The relative amounts of amplicons obtained for each of the four gene groups are depicted with the amounts of amplicons obtained with the bonobo genomic DNA as template being set to 1.0. Note that compared to chimpanzee and bonobo genomic DNA, the copy number in human genomic DNA is (i) two-fold higher for *ARHGAP11*, consistent with the presence of the human-specific gene *ARHGAP11B* in addition to the common gene *ARHGAP11A*; (ii) four-fold higher for *FAM72*, consistent with the presence of the human-specific genes *FAM72B, FAM72C* and *FAM72D* in addition to the common gene *FAM72A*; (iii) three-fold higher for *GTF2H2*, consistent with the presence of the human-specific genes *GTF2H2B* (black bold letters, not among the cNPC-enriched genes identified in this study) and *GTF2H2C* in addition to the common gene *GTF2H2A*; and (iv) two-fold higher for *SMN*, consistent with the presence of the human-specific gene *SMN2* in addition to the common gene *SMN1*.

The human-specific *NOTCH2NL* identified here as a cNPC-enriched gene arose from partial duplication of the *NOTCH2* gene (Fig. 4A) and comprises only those regions of the *NOTCH2* gene that give rise to a short *NOTCH2* splice variant (Ensembl transcript ENST00000602566.5). Similar to this short NOTCH2 isoform, *NOTCH2NL* encodes only a short segment of the NOTCH2 ectodomain. However, the protein encoded by the *NOTCH2NL* studied here is predicted to lack a signal peptide, which raises the issue of whether NOTCH2NL is secreted, and if so, via which pathway. Irrespective of this open question, we explored, in light of the importance of Notch signaling for cNPC behaviour (Imayoshi et al., 2013; Kawaguchi et al., 2008; Lui et al., 2011; Pierfelice et al., 2008; Wilkinson et al., 2013), a potential role of the *NOTCH2NL* mRNA and/or the NOTCH2NL protein in cNPCs by in utero electroproration of *NOTCH2NL* under the control of a constitutive promoter into neocortical aRG of embryonic day (E) 13.5 mouse embryos. Analysis by Ki67 immunofluorescence 48 hours after *NOTCH2NL* electroporation revealed an increase in cycling basal progenitors in the SVZ and IZ, but not in apical progenitors in the VZ (Fig. S1A, B). This finding was further corroborated by analysis of mitotic cNPCs using phosphohistone H3 immunofluorescence, which showed an increase in abventricular, but not ventricular, mitoses (Fig. S1C-E). Thus, forced expression of the human-specific *NOTCH2NL* gene in mouse embryonic neocortex appears to promote basal progenitor proliferation.

The remaining two human-specific cNPC-enriched protein-coding genes evolved in distinct ways. The *ZNF492* gene as such exists in the genomes of all non-human great apes. In the case of human, however, an exon of another zinc finger protein-encoding gene, *ZNF98*, inserted into the *ZNF492* locus, yielding a chimeric human-specific protein containing the repressor domain of ZNF492 and the DNA binding domain of ZNF98 (Fig. 4C). The *FAM182B* gene as such exists not only in human but also in chimpanzee, bonobo and gorilla. However, in bonobo and gorilla, a stop codon terminates the potential open reading frame soon after the initiator methionine, whereas in human a single T–>G substitution abolishes this stop codon and rescues the open reading frame to yield a 152-amino acid-long protein (Fig. 4D). In chimpanzee, the corresponding T is missing, resulting in a reading frame shift that predicts a shorter, 52-amino acid-long polypeptide. Taken together, we conclude that the human-specific cNPC-enriched protein-coding genes evolved by a variety of evolutionary mechanisms.

We sought to corroborate that the human-specific cNPC-enriched protein-coding genes arising from complete or partial gene duplication indeed constitute additional gene copies (rather than reflecting the inability of distinguishing multiple gene copies in the genomes of the other great apes due to genome assembly issues). To this end, we used a quantitative genomic PCR approach. The idea was that primers targeting genomic regions that are identical in human, chimpanzee and bonobo and thus should amplify genomic DNA of the three species proportionally to the copy number of each gene in each species. As a proof of principle, we validated the known human-specific nature of the partially duplicated *ARHGAP11B* by designing primers to the regions that are identical between *ARHGAP11A* and *ARHGAP11B*. Using the bonobo gene as the standard, this resulted in a two-fold increase of the human PCR product compared to the bonobo and chimpanzee, corroborating that *ARHGAP11B* is indeed a human-specific partial gene duplication (Fig. 4E).

We used the same approach to validate the human-specific genes arising from complete gene duplication. For four of these nine genes (*ANKRD20A2, ANKRD20A4, CBWD5, DHRS4L2*) and for *NOTCH2NL*, we could not design primers that uniquely target these genes as the respective genomic loci are not well resolved in the non-human great ape genomes. Thus, the final validation of these putative human-specific genes awaits improved genome assemblies. For the other five human-specific cNPC-enriched genes (*FAM72B/C/D, GTF2H2C, SMN2*) and for the human-specific gene *GTF2H2B* for which primers could be designed, genomic qPCR resulted in an estimated four human copies of *FAM72*, three human copies of *GTF2H2* and two human copies of *SMN* (Fig. 4E), compared to only one copy in both chimpanzee and bonobo. This validated the human-specific nature of these genes.

### Spatial mRNA expression analysis in fetal human neocortex of the human-specific cNPC-enriched protein-coding genes and of three selected primate-but not human-specific protein-coding genes

Given that the 13 human-specific genes had emerged from a screen for cNPC-enriched genes, it was of interest to examine their spatial expression pattern in the various zones of the fetal human cortical wall. To this end, we performed in-situ hybridization (ISH) on 13 wpc human neocortex to determine the localization of their mRNAs. Depending on the gene under study, this analysis detected expression of either the human-specific gene only, or (if the ISH probe used could not distinguish between paralogs due to their sequence similarity) the mRNA of the ancestral paralog from which the duplication arose and – if existing – the mRNAs of yet other paralogs. Of note, for *ARHGAP11B*, we used a specific Locked Nucleic Acid (LNA) probe, which enabled us to distinguish the mRNA of *ARHGAP11B* from that of *ARHGAP11A* (Fig. S2). We could distinguish five types of gene expression patterns.

First, referred to as "VZ", mRNA expression essentially confined to the VZ, which was the case for *ANKRD20A1-4* and *NOTCH2NL* (Fig. 5A-B"'). Second, referred to as "VZ + iSVZ + oSVZ", mRNA expression in all three germinal zones but not in the CP, which was the case for *ARHGAP11B* (Fig. 5C-C"'). Third, referred to as "VZ, iSVZ > oSVZ, CP", mRNA expression in all zones, however with markedly stronger staining in the VZ and iSVZ than in the oSVZ and CP, which was the case for *DHRS4L1-2, FAM72A-D* and *ZNF492* (Fig. 5D-F'"). Fourth, referred to as "VZ > CP > iSVZ, oSVZ", mRNA expression in all zones, however with markedly stronger staining in the VZ and CP than in the iSVZ and oSVZ, which was the case for *GTF2H2A-C* (Fig. 5J-J'"). Fifth, "referred to as VZ, CP > iSVZ, oSVZ", strong mRNA expression in the VZ and CP and lower mRNA expression in the iSVZ and oSVZ, which was the case for *CBWD1-7, FAM182A-B* and *SMN1-2* (Fig. 5G-I'").

**Fig. 5.**
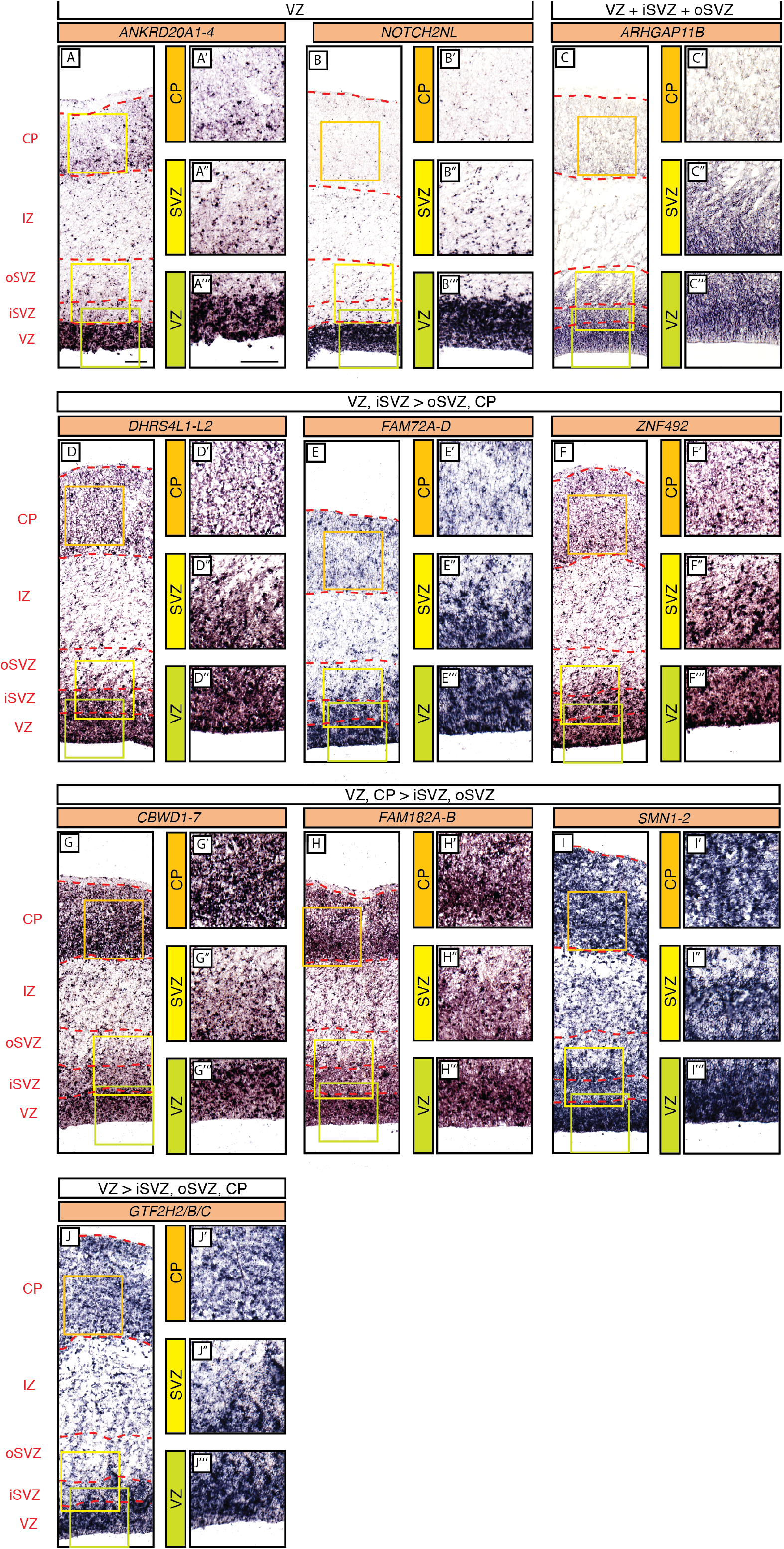
In-situ hybridization analysis of the mRNA levels of the human-specific cNPC-enriched protein-coding genes in the various zones of the fetal neocortical wall. Coronal sections of human fetal neocortex (13wpc) were subjected to ISH using probes either specific for the mRNA of the human-specific gene under study (B, C, F) or recognizing the mRNAs of both the human-specific gene(s) and the paralog gene common to other primates as well (A, D, E, G, H, I, J). The five patterns of preferential mRNA expression in the various zones of the fetal neocortical wall (see labeling on the left and red dashed lines) are indicated above the images. Green, yellow and orange boxes indicate areas of the VZ, SVZ and CP, respectively, that are shown at higher magnification in the respective images labeled "', " and ' on the right. Scale bars, 100 μm.

The strong ISH signal in the CP observed for the latter genes is nonetheless consistent with our conclusion, based on our approach of gene identification (Fig. 1), that the human-specific paralogs show enriched expression in cNPCs. Indeed, the published RNA-Seq data from specific LCM-isolated neocortical zones (Fietz et al., 2012) confirmed the ISH pattern for *CBWD1-7, FAM182A-B* and *SMN1-2* in that the sum of their mRNA levels in the three germinal zones (VZ, iSVZ, oSVZ) was greater than the mRNA level in the CP.

We also examined by ISH the spatial expression pattern in the fetal human cortical wall of the three primate-specific genes *PTTG2, MICA* and *KIF4B*. Due to the high degree of similarity in nucleotide sequence this analysis also included the mRNA of the respective ancestral paralog. mRNA expression for *PTTG1/2* (Fig. 6A), *MICA/B* (Fig. 6B) and *KIF4A/B* (Fig. 6C) was robust in the human VZ and iSVZ, relatively low in the oSVZ, and moderate in the CP.

**Fig. 6.**
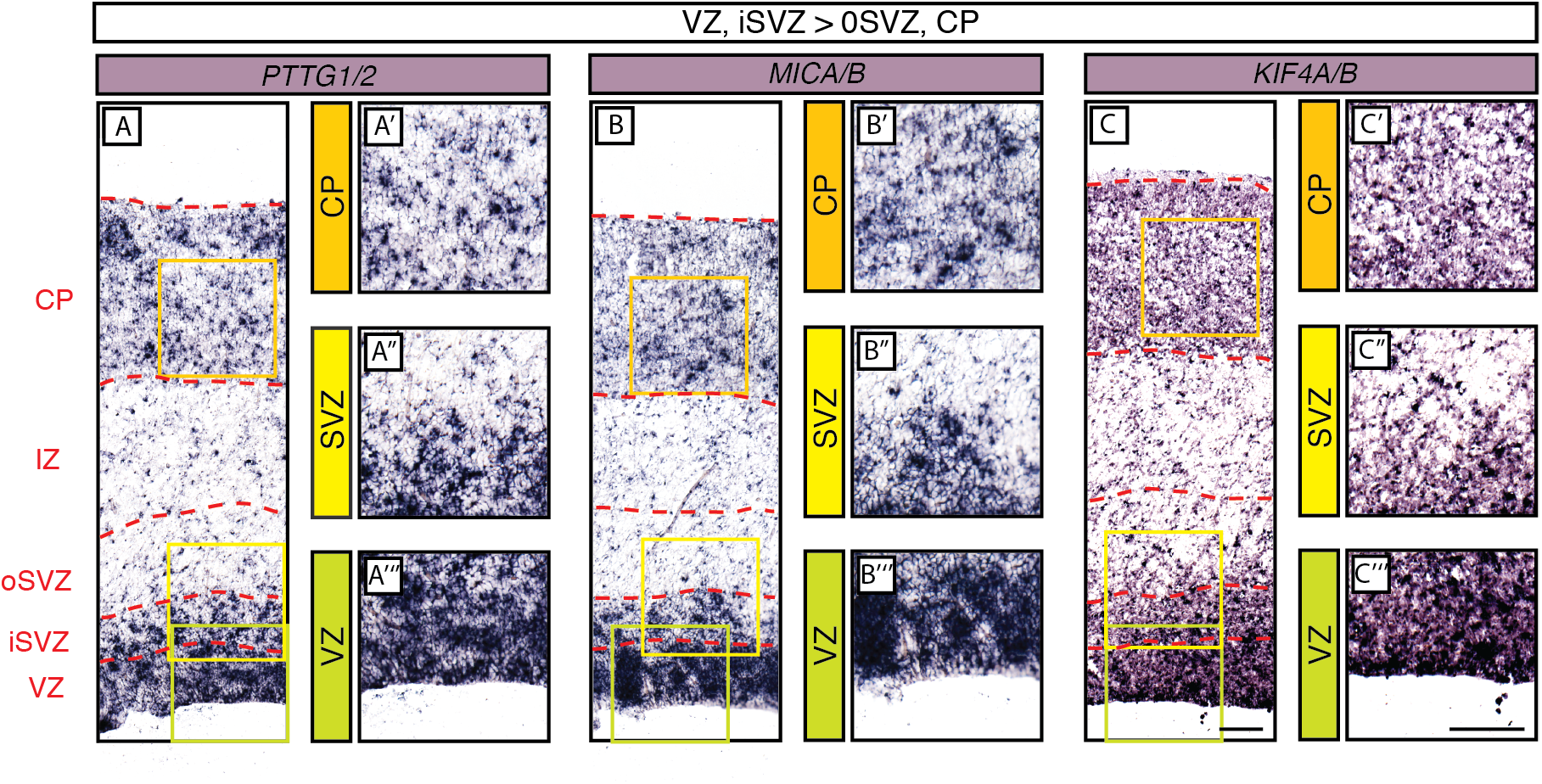
In-situ hybridization analysis of the mRNA levels of 3 selected primate-specific genes in the various zones of the fetal human neocortical wall. Coronal sections of human fetal neocortex (13wpc) were subjected to ISH using probes recognizing the mRNAs of either both, the human cNPC-enriched protein-coding gene under study that has orthologs specific to non-human primates (**A**, *PTTG2;* C, *KIF4B*) and the paralog gene common to non-primates as well (**A**, *PTTG1;* C, *KIF4A*), or the human cNPC-enriched protein-coding gene *MICA* that has orthologs specific to non-human primates and a paralog specific to primates, *MICB* (**B**). The pattern of preferential mRNA expression of the 3 genes under study in the various zones of the fetal neocortical wall (see labeling on the left and red dashed lines) is indicated above the images. Green, yellow and orange boxes indicate areas of the VZ, SVZ and CP, respectively, that are shown at higher magnification in the respective images labeled "', " and ' on the right. Scale bars, 100 μm.

### Cell type-specific expression patterns of the human-specific cNPC-enriched protein-coding genes compared to the corresponding ancestral paralogs

Complete or partial gene duplications often encompass the regulatory elements that control gene expression (Bailey et al., 2002; Eichler et al., 2004; Fortna et al., 2004; Hurles, 2004). This raises the question whether the human-specific cNPC-enriched protein-coding genes identified here exhibit similar cell-type expression patterns as their respective ancestral paralogs, or whether expression differences have evolved during human evolution.

Given that we could not distinguish by ISH most of the human-specific genes from their respective ancestral paralog, we sought an additional approach to gain insight into the cell-type specificity of expression of the human-specific genes. Specifically, we used our previously reported cell-type-specific gene expression data from the human aRG population (aRG), the bRG population (bRG) and the neuron fraction (N) (Florio et al., 2015) and re-analyzed these data using Kallisto. Kallisto is a probabilistic algorithm to estimate absolute transcript abundance, which has been proven to be accurate in assigning RNA-Seq reads to specific transcripts, including those originating from highly similar paralog genes (Bray et al., 2016). We could confidently ascertain cell-type-specific mRNA expression profiles for 11 of the 13 human-specific genes and their corresponding ancestral paralog (Fig. 7, Table S4).

**Fig. 7.**
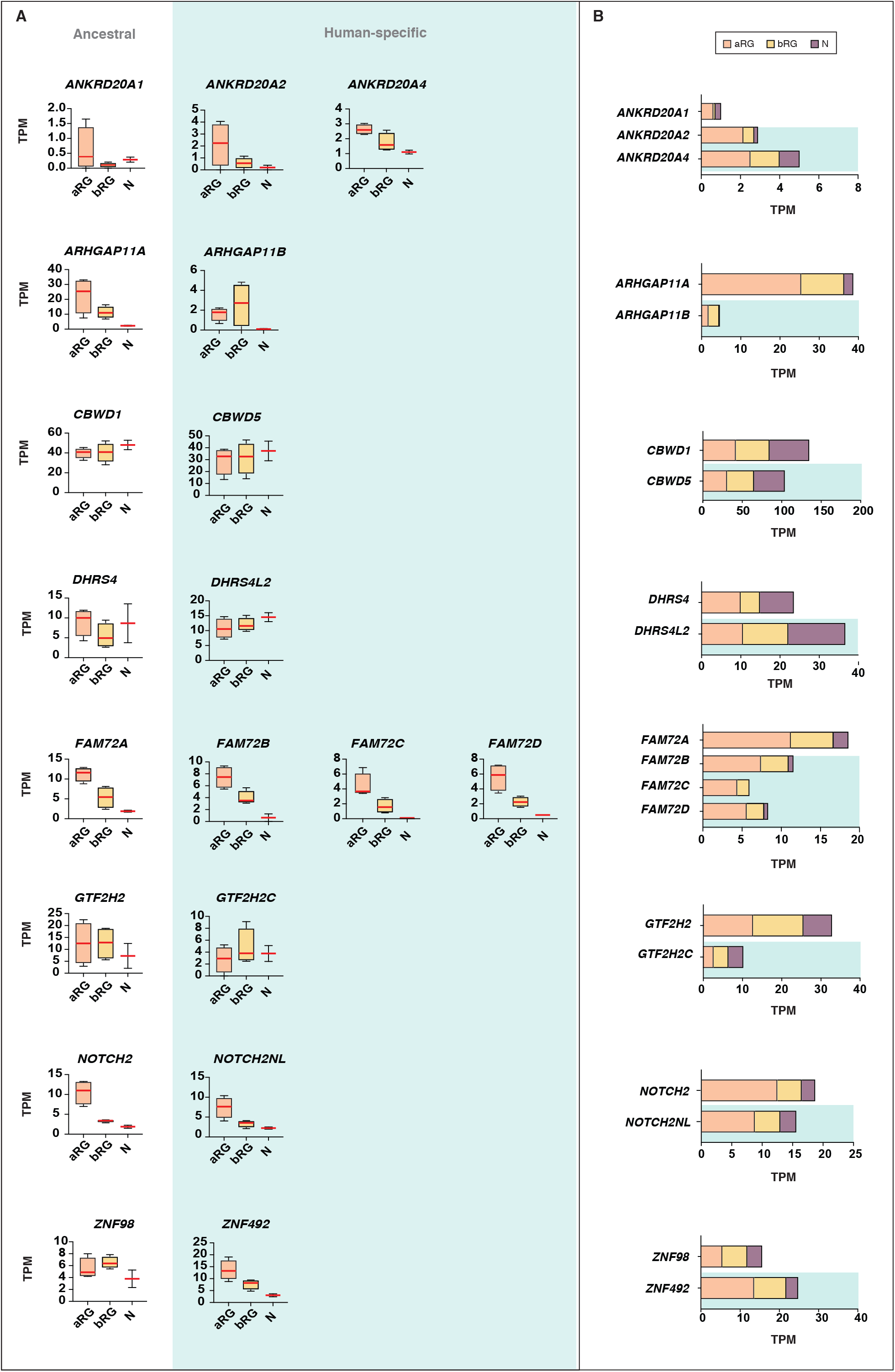
Comparison of the mRNA expression of 11 human-specific cNPC-enriched protein-coding genes with their ancestral paralogs in isolated cell populations enriched in aRG, bRG and neurons from fetal human neocortex. A previously published genome-wide transcriptome dataset obtained by RNA-Seq of cell populations isolated from fetal human neocortex, that is, aRG (orange) and bRG (yellow) in S-G2-M and a fraction enriched in neurons but also containing bRG in G1 (N, purple) (Florio et al., 2015), was analyzed for the abundance of mRNA-Seq reads assigned to either the indicated human-specific gene(s) under study (blue background) or the corresponding ancestral paralog (white background), using the Kallisto algorithm. (**A**) Min-max box-and-whiskers plots showing mRNA levels (expressed in Transcripts Per Million, TPM); red lines indicate the median. (**B**) Stacked bar plots showing the cumulative mRNA expression levels in the indicated cell types (sum of the median TPM values shown in (A)).

We first focused on changes in the total mRNA levels between the human-specific genes and their ancestral paralogs. With the exception of *CBWD5* and *NOTCH2NL*, for which the total mRNA levels in aRG, bRG and N were in the same range as the corresponding ancestral paralog, we found that the majority of the human-specific genes showed markedly different mRNA expression levels compared to their ancestral paralog, which were either reduced (*ARHGAP11B, FAM72B/C/D, GTF2H2C*) or increased (*ANKRD20A2, ANKRD20A4, DHRS4L2, ZNF492*) (Fig. 7A, B). This reflects either changes in mRNA expression levels per cell, changes in the proportions of mRNA-expressing cells, or both. Irrespective of which is the case, this finding indicates that expression of these human-specific genes has indeed changed compared to their ancestral paralogs during human evolution – a potential indication of neofunctionalization.

Next, we asked whether the human-specific genes diverged in their pattern of expression in aRG vs. bRG vs. N from that of their ancestral paralogs. For five of the human-specific genes (*CBWD5, FAM72B/C/D* and *NOTCH2NL*), the pattern of relative mRNA levels in these cell types was similar to that of the respective ancestral paralog (Fig. 7A). In the case of the other six human-specific genes, we observed, relative to the mRNA level in aRG, either decreases (*ZNF492*) or increases (*ARHGAP11B, DHRS4L2, GTF2H2C*) in the bRG mRNA level compared to respective ancestral paralog. We also observed decreases (*ANKRD20A2, ANKRD20A4*) or increases (*DHRS4L2, GTF2H2C*) in the N fraction mRNA level compared to the respective ancestral paralog (Fig. 7A). Of note, the increase in the *ARHGAP11B* mRNA level in bRG as compared to aRG is consistent with the previously reported function of this gene in basal progenitor amplification (Florio et al., 2015; Florio et al., 2016). These findings suggest that these six human-specific genes underwent changes in regulatory elements at the transcriptional and/or post-transcriptional level.

To complement these data, we performed a second type of analysis. We identified paralog-specific sequencing reads (Suppl. Files S1-S8; see Suppl. Fig. S3A for illustration of a hypothetical example) using our previously reported RNA-Seq dataset (Florio et al., 2015), and then determined the number of paralog-specific sequencing reads for the 11 human-specific genes and their corresponding ancestral paralog in aRG, bRG and N (Suppl. Fig. S3B). This analysis largely corroborated the results shown in Fig. 7A, further pointing to expression changes in aRG vs. bRG vs. N for the human-specific genes in comparison to their corresponding ancestral paralogs.

We finally explored the complexity in cell-type-specific expression patterns by examining the differential mRNA expression of protein-coding splice variants of the human-specific genes. Specifically, we analyzed our aRG vs. bRG vs. N RNA-Seq data (Florio et al., 2015) for cell-type-specific gene expression and relative abundance of sequencing reads diagnostic of specific protein-coding splice variants of 10 of the 11 human-specific cNPC-enriched genes shown in Fig. 7 (Suppl. Fig. S4). This showed, for most of these human-specific genes (*ANKRD20A2, ANKRD20A4, CBWD5, FAM72B/C, DHRS4L2, GTF2H2C, NOTCH2NL*), the preferential expression of certain splice variants. Moreover, this analysis revealed splice variants with preferential expression in either aRG, bRG or N for some of these human-specific genes (e.g., *CBWD5, GTF2H2C, ARHGAP11B*). A notable case was *ARHGAP11B*, of which one splice variant (Ensembl transcript ENST00000428041.2), endowed with a shorter 3'-UTR, was exclusively expressed in bRG whereas the other splice variant was enriched in aRG (Suppl. Fig. S4).

In summary, these analyses show that after duplication, the expression pattern of most of the resulting new, human-specific cNPC-enriched protein-coding genes evolved differences in both the levels and cell-type specificity of their mRNAs compared to their respective ancestral paralog.

## Discussion

Our study not only provides a resource of genes that are candidates to exert specific roles in the development and evolution of the primate, and notably human, neocortex, but also has implications regarding (i) the emergence of these genes during primate evolution and (ii) the maintenance vs. modification of the cell-type specificity of their expression. As to their emergence during primate evolution, two aspects of our findings deserve comment. First, while entire or partial gene duplications were the underlying mechanism that gave rise to the majority of the human-specific cNPC-enriched protein-coding genes, as noted previously (Bailey et al., 2002; Eichler et al., 2004; Fortna et al., 2004; Hurles, 2004), our data reveal also other mechanisms of gene evolution such as exon duplication and replacement (*ZNF492*) and translational stop codon removal (*FAM182B*).

The latter notion is further underscored by our observation that of the three primate-but not human-specific genes studied here in greater detail, two (*KIF4B, PTTG2*) arose by retroposition (Brosius, 1991; Long et al., 2003; Marques et al., 2005) rather than gene duplication. Here, *PTTG2* is a particularly interesting case in that its initially presumably open reading frame became closed during the evolution of non-hominoidea simiiformes but remained open during the evolution of hominoidea. This suggests that the functional role of PTTG2 may be essential for the development of the neocortex of apes and human but not for that of New-World and Old-World monkeys. Given the expression of *PTTG2* in the germinal zones of fetal human neocortex and the fact that this gene is derived from *PTTG1*, which encodes a protein exhibiting tumorigenic activity (Vlotides et al., 2007), it appears possible that PTTG2 may function to amplify cNPCs.

Second, of the 54 human genes that we identified in the present study as being primate-specific, as many as 13 (i.e. almost one quarter) are human-specific. This is a far greater percentage than would be expected if the former genes arose by a constant rate during primate evolution to modern humans. This in turn suggests that the latter cNPC-enriched protein-coding genes conveyed a selection advantage specifically during the evolution of the human neocortex.

As to the issue of maintenance vs. modification of the cell-type specificity of expression of the human-specific genes, it is striking to observe that the majority of these genes, although arising by entire or partial gene duplications, show marked differences not only in the level but also in the cNPC-type specificity of their mRNA expression compared to their ancestral paralog. For several of the human-specific genes, the corresponding spatial characteristics of their mRNA expression in the neocortical germinal zones could be corroborated by specific ISH. These data suggest that during human evolution these genes underwent specific changes in regulatory elements at the transcriptional and/or post-transcriptional level. This in turn raises the possibility that (at least some of) the human-specific genes characterized in the present study may be candidates to have contributed to the evolution of human-specific features of neocortical development.

In line with the latter consideration, we found that expressing the human-specific gene *NOTCH2NL* studied here in mouse embryonic neocortex increased the abudance of cycling basal progenitors, a hallmark of the developing human neocortex. Moreover, we previously showed that the human-specific function of *ARHGAP11B* in cNPCs arose by a single nucleotide substitution that generated a new splice donor site, the use of which generates a novel human-specific C-terminal protein sequence that we implicate in basal progenitor amplification (Florio et al., 2015; Florio et al., 2016). Importantly, this single nucleotide substitution presumably occurred relatively recently during human evolution (Florio et al., 2016), that is, after the partial gene duplication event ~5 million years ago (Antonacci et al., 2014; Dennis et al., 2017; Riley et al., 2002). Furthermore, we have identified here an *ARHGAP11B* splice variant that is specifically expressed in human bRG (Fig. S4), the basal progenitor type thought to have a key role in neocortex expansion (Betizeau et al., 2013; Borrell and Götz, 2014; Borrell and Reillo, 2012; Florio and Huttner, 2014; Lui et al., 2011). Interestingly, in contrast to the other protein-coding *ARHGAP11B* splice variant detected, which contains a long 3'-UTR with predicted microRNA binding sites and which is predominantly expressed in aRG, the bRG-specific *ARHGAP11B* splice variant contains only a short 3'-UTR lacking predictable microRNA binding sites. This suggests that *ARHGAP11B* mRNAs may be subject to differential, microRNA-mediated, regulation depending on whether ARHGAP11B functions in the lineage progression from aRG to bRG or in bRG amplification. Taken together, our findings reveal genomic changes at a variety of levels that gave rise to novel functions and patterns of expression in cNPCs and that are likely relevant for the development and evolution of the human neocortex.

## Materials and Methods

### Human fetal brain tissue

Human fetal brain tissue was obtained from the Klinik und Poliklinik für Frauenheilkunde und Geburtshilfe, Universitätsklinikum Carl Gustav Carus of the Technische Universität Dresden, following elective termination of pregnancy and informed written maternal consent, and with approval of the local University Hospital Ethical Review Committees. The gestational age of the specimen used for ISH (13 weeks post conception, wpc) was assessed by ultrasound measurements of crown-rump length, as described previously (Florio et al., 2015). Immediately after termination of pregnancy, the tissue was placed on ice and transported to the lab. The sample was then transferred to ice-cold Tyrode’s solution, and tissue fragments of cerebral cortex were identified and dissected. Tissue was fixed in 4% paraformaldehyde in 120 mM phosphate buffer (pH 7.4) for 3 hours at room temperature followed by 24 hours at 4°C. Fixed tissue was then incubated in 30% sucrose overnight, embedded in Tissue-Tek OCT (Sakura), and frozen on dry ice. Cryosections of 12 μm were produced using a cryostat (Microm HM 560, Thermo Fisher Scientific) and stored at −20°C until processed for ISH.

### Identification of human cNPC-enriched protein-coding genes

To identify genes the expression of which is enriched in human cNPCs, we screened differential gene expression data from four published datasets (Fietz et al., 2012; Florio et al., 2015; Miller et al., 2014; Pollen et al., 2015) generated from 12-19 wpc human fetal neocortex, using diverse cortical zone or cell type-enrichment strategies and modes of determination of RNA levels (summarized in Table S1).

<u>Fietz et al., 2012</u> – This dataset was generated by RNA-Seq of the germinal zones (VZ, iSVZ, oSVZ) and CP isolated by LCM from the neocortex of six human fetuses ranging in gestational age from 12 to 16 wpc. We screened this dataset for protein-coding genes more highly expressed, across all stages, in either VZ, iSVZ or oSVZ than CP (as determined by DGE analysis, p<0.01, (Fietz et al., 2012). The resulting data-subset contained 2,758 genes (Table S1, Fig. 1).

<u>Miller et al., 2014</u> (BrainSpan Atlas of the Allen Brain Institute, Prenatal LMD Microarray, http://www.brainspan.org/lcm/search/index.html) – This dataset (Miller et al., 2014) was generated by microarray RNA expression profiling of germinal zones (VZ, iSVZ, oSVZ) and neuron-enriched layers (IZ, subplate, CP, marginal zone, subpial granular zone) isolated by LCM from fetal human neocortex (for the purpose of the present analysis, only data obtained from two 15-16 wpc human fetuses were considered). We screened this dataset for protein-coding genes with highest correlation with either VZ, iSVZ or oSVZ (correlation coefficient >0.5) compared to all cortical regions analyzed. The resulting data-subset contained 4,555 genes (Table S1, Fig. 1).

<u>Florio et al., 2015</u> – This dataset was generated by RNA-Seq of human radial glia subtypes (aRG and bRG) and CP neurons (N) isolated from the neocortex of two 13 wpc human fetuses. These cell types were differentially labeled using a combination of fluorescent molecular markers, and isolated by FACS. By experimental design, only cells that exhibited apical plasma membrane and/or contacted the basal lamina were isolated. Moreover, the isolation of aRG and bRG was confined to cells that had duplicated their DNA, and the neuron fraction contained a minority of bRG in G1 (Florio et al., 2015). We screened this dataset for protein-coding genes with higher expression in either aRG or bRG than N (as determined by DGE analysis, p<0.01, (Florio et al., 2015). The resulting data-subset contained 2,106 genes (Table S1, Fig. 1).

<u>Pollen et al., 2015</u> – This dataset was generated by RNA-seq of single cells captured from the VZ and SVZ microdissected from the neocortex of three 16.5-19 wpc human fetuses. Cells were post-hoc attributed – based on gene expression profiling – to either radial glia (aRG and bRG), intermediate progenitors (i.e. bIPs), or neurons (N). We screened this dataset for genes positively correlated with either radial glia or bIPs (correlation coefficient >0.1, (Pollen et al., 2015) and negatively correlated with N (correlation coefficient <0.1, (Pollen et al., 2015). The resulting data-subset contained 5,335 genes (Table S1, Fig 1).

These data-subsets contain only protein-coding genes, which were identified and selected using the Ensembl data-mining tool BioMart (http://www.ensembl.org/biomart/martview/), implementing the Genome Reference Consortium Human Build 38 (GRCh38.p10) dataset.

Next, we intersected the four data-subsets obtained. To do this, we converted all gene IDs contained in the four original datasets to match the latest Ensembl gene annotation (Ensembl v89) of the GRCh38.p10 genome assembly, and then searched for the co-occurrence of genes (or lack thereof) across the four data-subsets. This resulted in 3,722 human cNPC-enriched protein-coding genes present in at least two of the four data-subsets (listed in Table S1, see also Fig. 1).

### Screening of human cNPC-enriched protein-coding genes for primate-specific orthologs

The 3,722 human cNPC-enriched protein-coding genes were screened for the occurrence of one-to-one orthologs in non-primate species, using BioMart and implementing v89 Ensembl annotation of “1-to-1 orthologs”. All genes that had an annotated 1-to-1 ortholog in non-primate species were excluded from our four data-subsets. This yielded 83 genes that were candidates to be primate-specific.

We visualized whole genome alignments in the UCSC genome browser (Tyner et al., 2017) to manually analyze each of the 83 candidate primate-specific genes. To this end, we inspected co-linear chains of local alignments (Kent et al., 2003) between the human hg38 genome assembly and the assemblies of non-primate mammals to check if the human gene locus aligned to non-primate mammals. For the genes that aligned to non-primate mammals, regardless of whether they aligned in a conserved or in a different context, we used gene annotations of the aligning species to assess which gene is annotated in the respective locus. For this purpose, we made use of gene annotations from Refseq, Ensembl (Aken et al., 2017) and CESAR (Sharma et al., 2016) (a method that transfers human gene annotations to other aligned genomes if the gene has an intact reading frame), and removed those candidate genes that likely have an aligning gene in non-primate mammals. This reduced the list of the 83 candidates to 54 genes that we considered as primate-specific.

### Tracing the evolution of the primate-specific genes in the primate lineage

We traced the evolution of these 54 primate-specific genes in the primate lineage to determine which of these have orthologs, in non-human primates, to the corresponding 54 human cNPC-enriched protein-coding genes, and which do not, and therefore are human-specific. To this end, we inspected co-linear alignment chains and a multiple genome alignment that includes 17 non-human primate genomes (Sharma and Hiller, 2017). For the genes that aligned to other primates, we used the CESAR annotations to check if a gene of interest has an intact reading frame in other species. We only considered a gene to be conserved if an intact reading frame is present in the respective species. For example, while *FAM182* aligns in a conserved context to chimpanzee and gorilla, CESAR did not find an intact reading frame and did not annotate the gene; indeed, inspecting the multiple genome alignment revealed a frameshift in chimpanzee and a stop codon mutation in gorilla, showing that *FAM182B* is likely a non-coding gene in non-human primates. Then, we assigned each gene to a node in the primate phylogeny (clade), based on the descending species that likely have an intact coding gene. Note that this inferred ancestry does not imply that all descending species have an intact gene. This is exemplified by *TMEM99*, which aligns to all great apes and has an intact reading frame in human and orangutan, but encodes no or a truncated protein in chimpanzee/bonobo (due to a frameshift mutation) and gorilla (due to a stop codon mutation).

We combined this analysis with BLAT searches using the human protein or human mRNA sequence to assess the number of aligning loci in other primates; however, this was not conclusive for highly complex loci such as the duplications involving *ANKRD20A* and *CBWD5* genes, where numerous similar genes and pseudogenes are present and the completeness of non-human primate genome assemblies is not certain due to the presence of assembly gaps. In addition, for human-specific candidates that arose by duplication, inspecting the respective genomic locus in the chimpanzee genome browser was useful, since human duplications are visible as additional, overlapping alignment chains.

### Paralog-specific and isoform-specific gene expression

To estimate expression differences among cNPC types between (a) given human-specific gene(s) and its/their highly similar ancestral paralog(s) in the human genome, we used the Kallisto probabilistic algorithm, which has been proven to be accurate in assigning reads to specific transcripts, including those originating from highly similar paralog genes in the human genome (Bray et al., 2016).

For this analysis, we used reads generated previously by RNA-Seq of human aRG, bRG and N (SRA Access, SRP052294, (Florio et al., 2015)) as input, GRCh38 as genome reference, and Ensembl v89 as genome annotation reference. Transcript abundances were output in Transcripts per Million (TPM) units. To compare expression between human-specific and ancestral paralog genes (Fig. 7), we extracted TPM values for all paralogs in each orthologous group, and summed the TPM values for all protein-coding transcripts (as per Ensembl annotation) for each gene. To compare expression between different splice variants produced by each human-specific gene (Fig S4), we extracted the TPM values specific for each individual splice variant and expressed the data relative to each other.

Kallisto’s transcript abundance measurements represent a probabilistic approximation of actual transcript levels, and thus are an estimate. In order to compare actual paralog gene expression in distinct cNPC types and neurons, we performed a second type of analysis, which did not aim at providing an estimate of absolute transcript abundances, but rather at providing a precise determination of the relative gene expression differences between paralogs. To this end, we aligned mRNA sequences of ancestral and human-specific paralogs in each orthology group, using Clustalw2 (http://www.ebi.ac.uk/Tools/msa/clustalw2/), and manually identified the homologous (but not identical) core sequence of each alignment (Suppl. Files S1-S8; see Fig. S3A for illustration of a hypothetical example). The corresponding sequences of each paralog – of same length by design – were used as reference for previously generated RNA-Seq reads from aRG, bRG and N (SRA Access, SRP052294, (Florio et al., 2015)) in order to search for paralog-specific mRNA reads. Reads aligning to both, ancestral and human-specific paralogs, were discarded as ambiguous, and only those reads aligning to paralog-specific sites (SNPs or indels), referred to as paralog-specific reads, were used for quantification (Fig S3B). This stringent alignment was carried out using bowtie1 (bowtie -Sp 5 -m 1 -v0).

It should be noted that, in contrast to the Kallisto-based analysis, the latter type of analysis does not distinguish between reads that originate from protein-coding and non-protein-coding transcripts of a given gene. Therefore, the quantifications shown in Fig. S3B reflect counts of all reads mapping to a given gene, whereas the quantifications shown in Fig. 4 reflect summed counts of protein-coding gene transcripts only.

### Genomic qPCR

Genomic DNA was obtained from EBV-transformed B cells of human, bonobo and chimpanzee, as described previously (Prüfer et al., 2012). Primers (Table S2) were designed for two different amplicons per orthologous gene group to bind to the same region of the human-specific gene(s) under study, its human paralog(s), and the chimpanzee and bonobo orthologs. Only one mismatch in the primer binding sequence between the reference genomes of the three species was allowed.

qPCR was performed on human, chimpanzee and bonobo genomic DNA, using either the ABsolute qPCR SYBR Greenmix (Thermo Fisher Scientific) on a Mx3000P qPCR System (Stratagene) or the Fast Start Essential DNA Green Master (Roche) on a Lightcycler 96 (Roche). The relative copy number between the three species was determined by the comparative cycle threshold (Ct) approach (Livak and Schmittgen, 2001) as follows. The Ct values for the human, chimpanzee and bonobo genes under study were normalized to the Ct value of the highly conserved single-copy gene *STX12*. The normalized values were then compared between the three species, using bonobo as reference, to determine the relative copy number.

### In utero electroporation and tissue processing

In utero electroporation was performed on C57BL/6J mice in agreement with German Animal Welfare Legislation, as described previously (Florio et al., 2015). Pregnant dams carrying E13.5 embryos were deeply anesthetized using isoflurane. Embryos were injected into the lateral ventricle with either 1 μg/μl of pCAGGS-NOTCH2NL and 0.5 μg/μl of pCAGGS-GFP or 1 μg/μl of empty pCAGGS and 0.5 μg/μl of pCAGGS-GFP in PBS containing 0.1% Fast Green, followed by electroporation (30 V, six 50-msec pulses with 1 sec intervals). Electroporated cerebral cortices were dissected at E15.5 and fixed overnight at 4°C in 4% paraformaldehyde in 120 mM phosphate buffer (pH 7.4). Fixed cortices were incubated in 20% sucrose for 24 hours at 4°C. Cortices were embedded in Tissue-TEK (O.C.T, Sakura Finetek) and stored at −20°C.

### Immunofluorescence

Cryosections of 20 μm were prepared. Cryosections were first rehydrated in PBS. Antigen retrieval was performed for 1 hour at 70°C in 0.01 M citrate. Cryosections were permeabilized by treatment with 0.1% Triton X-100 in PBS for 30 min. Cryosections were quenched for 30 min in 0.1 M glycine in PBS, blocked in 0.2% gelatin, 300 mM NaCl, and 0.3% Triton X-100 in PBS, and incubated overnight at 4°C with primary antibodies (Ki67, rabbit, Abcam, Ab15580, 1/500; PH3, rat, Abcam, Ab10543, 1/1000; GFP, chicken, Abcam, Ab13970, 1/1000). Appropriate secondary antibodies were incubated for 2 hours at room temperature (Alexa Fluor 488, 594, Molecular Probes, 1:500; DAPI, Sigma, 1/1000). Cryosections were mounted in Mowiol (Merck Biosciences).

### In-situ hybridization

Templates were amplified by PCR (see Table S3 for primer sequences) from oligo-dT-primed cDNA prepared from fetal human neocortex total RNA, and RNA probes directed against the mRNA(s) of a given human-specific gene and (if applicable) its paralog(s) in the human genome were synthesized using the DIG RNA labeling Mix (Roche). The *ARHGAP11B* LNA probe was designed with the Custom LNA mRNA Detection Probe design tool (Exiqon), focusing only on the sequence spanning the *ARHGAP11B* exon5–exon6 boundary, where *ARHGAP11B* is sufficiently different from *ARHGAP11A* (see Fig. S2) (Florio et al., 2016), and searching for hybridization with a predicted RNA melting temperature of 85°C. The LNA probe (5'-AGTCTGGTACACGCCCTTCTTTTCT-3') was synthesized and labelled with digoxigenin at the 5’ and 3’ ends (Exiqon).

In-situ hybridization was performed on 12–μm cryosections of 13 wpc fetal human neocortex and on COS-7 cells. Prior to the hybridization step, cryosections/cells were sequentially treated with 0.2 M HCl (2x 5 min, room temperature) and then with 6 μg/ml proteinase K in PBS, pH 7.4 (20 min, room temperature). Hybridization was performed overnight at 65°C with either 20 ng/μl of a given RNA probe or 40 nM *ARHGAP11B* LNA probe. TSA Plus DIG detection Kit (Perkin Elmer) was used for signal amplification, and the signal was detected immunohistochemically with mouse anti-digoxigenin HRP antibody (Perkin Elmer) and NBT/BCIP (Roche) as color substrate.

### Image acquisition

ISH images were acquired on a Zeiss Axio Scan slide scanner, and processed using ImageJ. Fluorescent images of electroporated neocortex were acquired using a Zeiss laser scanning confocal microscope 700 using a 20x objective. Quantifications were performed using Fiji.

## Acknowledgements

We are grateful to the Computer Service Facilities of the MPI-CBG and MPI-PKS and to other services and facilities of the MPI-CBG for the outstanding support provided, notably J. Helppi and his team from the Animal Facility and Jan Peychl and his team from the Light Microscopy Facility. We thank Robert Lachmann for providing fetal human tissue, Hella Hartmann of the CRTD of the Technische Universität Dresden for help with the acquisition of the ISH images, and members of the Huttner laboratory for critical discussion. We are grateful to Dr. Tomislav Maricic (MPI for Evolutionary Anthropology) for donating human, chimpanzee and bonobo genomic DNA. M.F. would like to thank Dr. Fenna Krienen (Harvard Medical School) for helpful discussion and critical reading of the manuscript. M.F. was a member of the International Max Planck Research School for Cell, Developmental and Systems Biology and a doctoral student at Technische Universität Dresden. W.B.H. was supported by grants from the Deutsche Forschungsgemeinschaft (DFG) (SFB 655, A2), the European Research Council (250197), and ERA-NET NEURON (MicroKin).

## Figure Legends

**Fig. S1.**
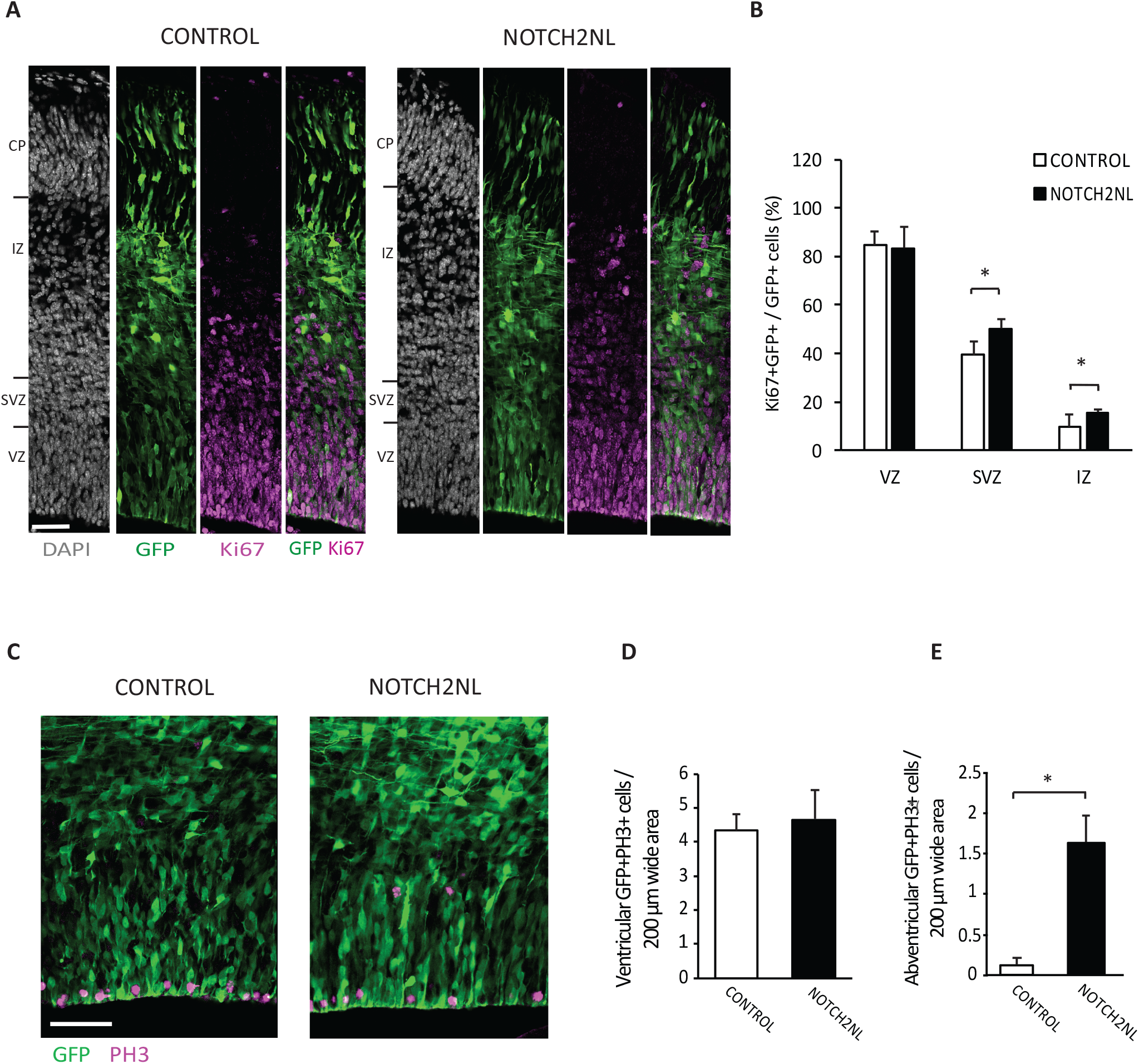
Forced expression of *NOTCH2NL* in mouse embryonic neocortex increases cycling basal progenitors. The neocortex of E13.5 mouse embryos was in utero co-electroporated with a plasmid encoding GFP together with either an empty vector (Control) or a *NOTCH2NL* expression plasmid (NOTCH2NL), all under constitutive promoters, followed by analysis 48 hours later. (**A**) GFP (green) and Ki67 (magenta) double immunofluorescence combined with DAPI staining (grey) of control (left) and NOTCH2NL-electroporated (right) neocortex. (**B**) Quantification of the percentage of targeted, i.e. GFP+, cells that are Ki67+ in the VZ, SVZ and IZ upon control (white bars) and *NOTCH2NL* (black bars) electroporation. (**C**) GFP (green) and phosphohistone H3 (PH3) double immunofluorescence of control (left) and NOTCH2NL-electroporated (right) neocortex. (**D, E**) Quantification of the number of ventricular (D) and abventricular (E) targeted (GFP+) cells in mitosis (PH3+) in a 200 μm-wide microscopic field upon control (white bars) and *NOTCH2NL* (black bars) electroporation. (**A, C**) Images are a single 2-μm optical sections. Scale bars, 50 μm. (**B, D, E**) Data are the mean of 10 embryos each, with 2-4 cryosections (**B**, 100 μm-wide microscopic field) per embryo counted and averaged. Error bars indicate SEM; *, *P* <0.05; Student’s *t*-test in B (VZ: *P* = 0.914, df = 18, *t* = 0.108; SVZ: *P* = 0.012, df = 18, *t* = 2.731; IZ: *P* = 0.020, df = 18, *t* = 2.501), **D** (*P* = 0.718, df = 18, *t* = 0.3656) and **E** (*P* = 0.011, df = 18 *t* = 2.841).

**Fig. S2.**
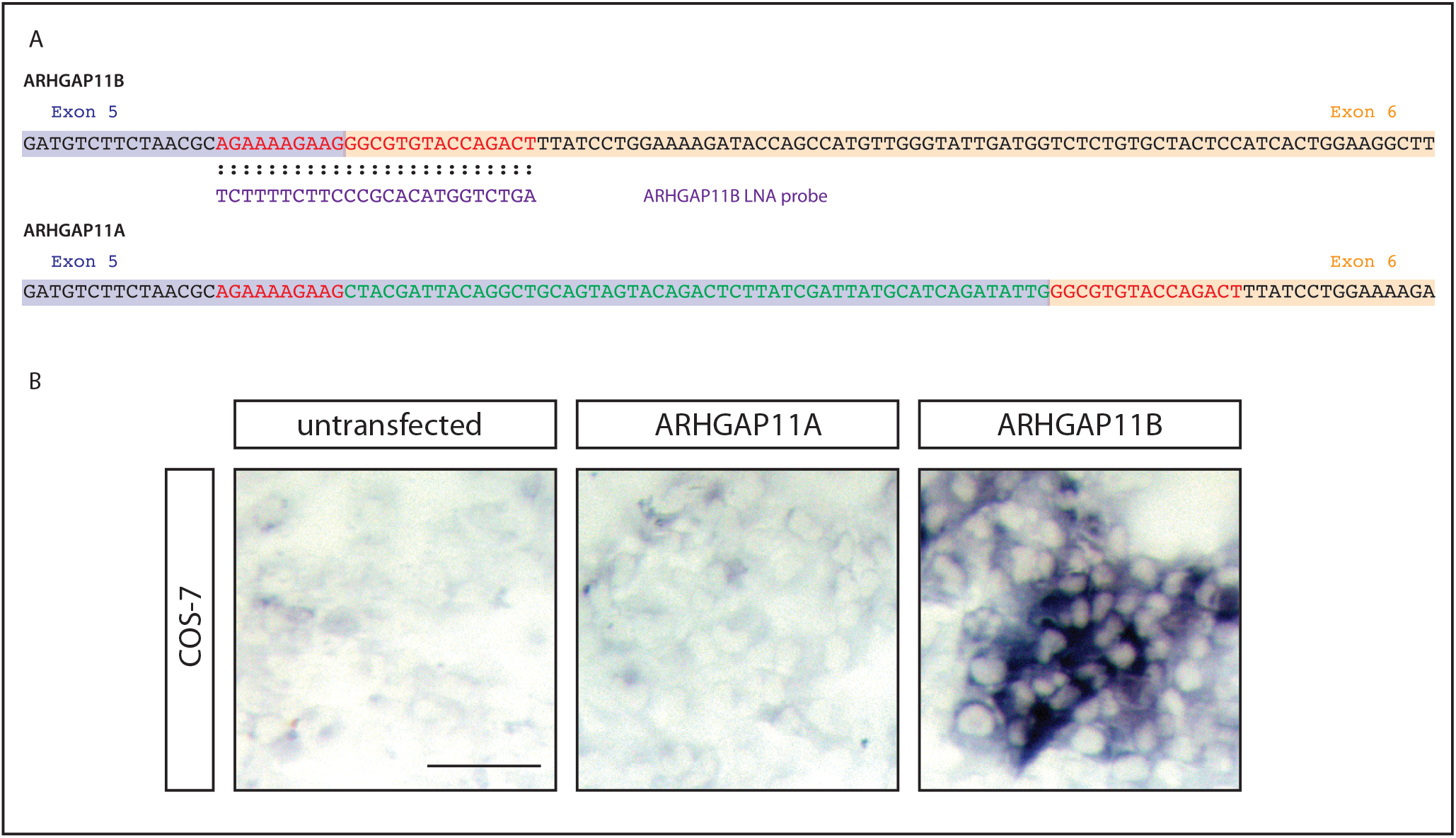
*ARHGAP11B-specific* ISH probe. (**A**) Nucleotide sequences at the exon 5 (purple background) – exon 6 (orange background) junction of the *ARHGAP11B* (top) and *ARHGAP11A* (bottom) mRNAs (note that U is depicted as T). The *ARHGAP11B* LNA ISH probe shown is are complementary to the nucleotides shown in red. The 55 nucleotides shown in green are unique to the 3'-end of the *ARHGAP11A* exon 5 and interfere with the binding of the LNA ISH probe to the *ARHGAP11A* mRNA, rendering the probe *ARHGAP11B-specific*. (**B**) Images of COS-7 cells that were either untransfected, or transfected with either an *ARHGAP11A-* or *ARHGAP11B-expressing* construct and stained with the *ARHGAP11B* LNA ISH probe. Note that an ISH signal is detected only in ARHGAP11B-transfected COS-7 cells, confirming the specificity of the LNA ISH probe for *ARHGAP11B*. Scale bar, 50 μm.

**Fig. S3.**
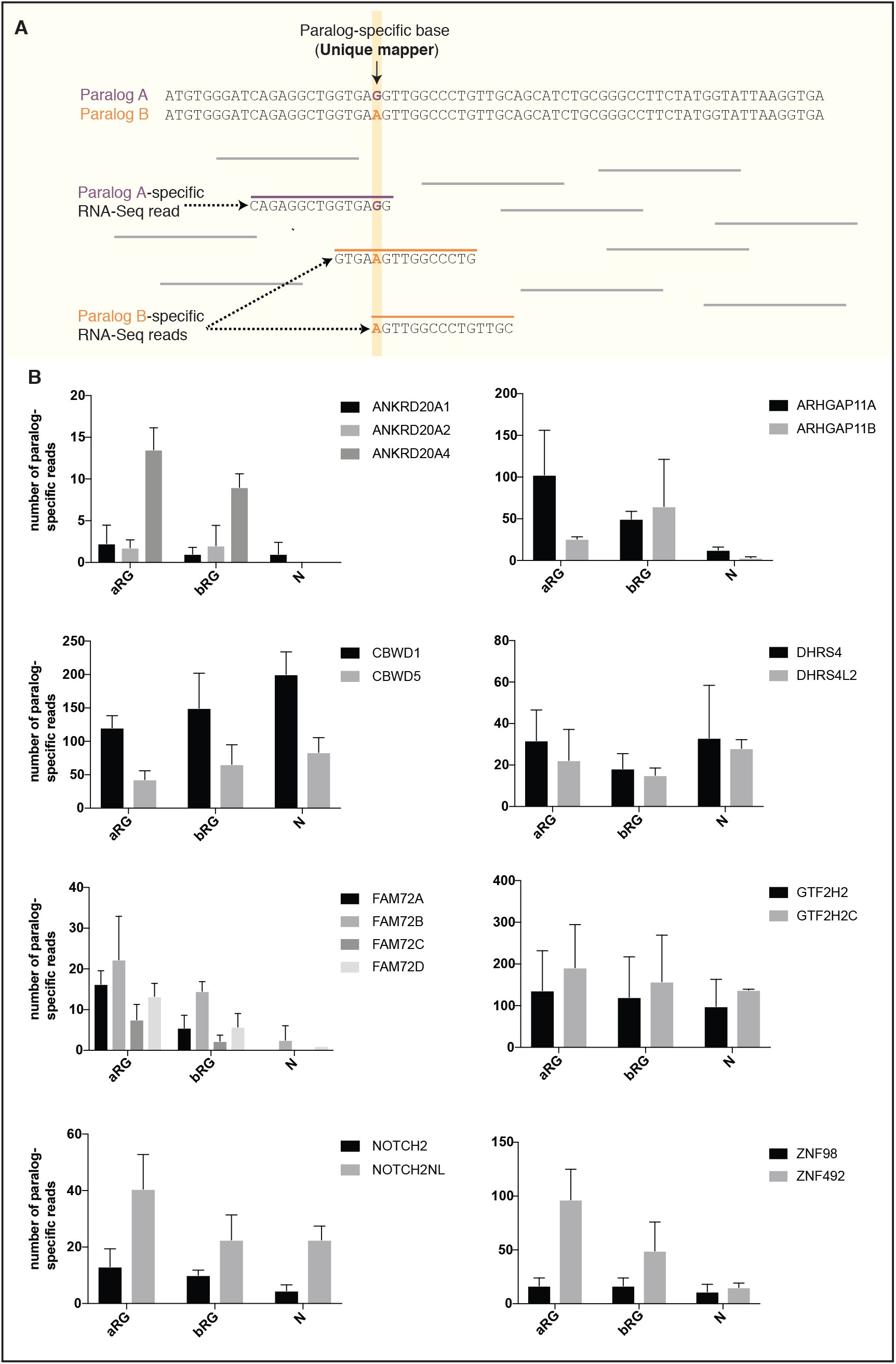
Comparison of the paralog-specific mRNA expression between 11 human-specific cNPC-enriched genes and their respective ancestral paralog in aRG, bRG and neuron-enriched cell populations from fetal human neocortex. (**A**) Diagram outlining the strategy used to ascertain paralog-specific mRNA expression in a given cell type of interest. mRNA sequences of an ancestral vs. a human-specific paralog (paralog A vs. B in the example shown) were aligned, and the homologous, yet distinct, core sequences of each alignment were extracted. The corresponding sequences of each paralog were used as a mapping reference for RNA-Seq reads from aRG, bRG and neuron-enriched cell populations from fetal human neocortex (Florio et al., 2015). Only reads aligning to “unique mappers”, i.e. paralog-specific sites (SNPs or indels), were used for the analysis shown in (B). In the example shown, paralog-specific reads specific for paralog A or paralog B, as defined by the paralog-specific base (vertical yellow line) are colored in purple and orange, respectively. (**B**) Bar plots showing the total numbers of paralog-specific RNA-Seq reads (identified as described in (A)) found in aRG vs. bRG vs. neuron-enriched (N) cell populations from fetal human neocortex (Florio et al., 2015). Grey bars indicate human-specific genes; black bars indicate their respective ancestral paralog. Data are the mean of four individual samples isolated from two human specimens; errors bars, SD.

**Fig. S4.**
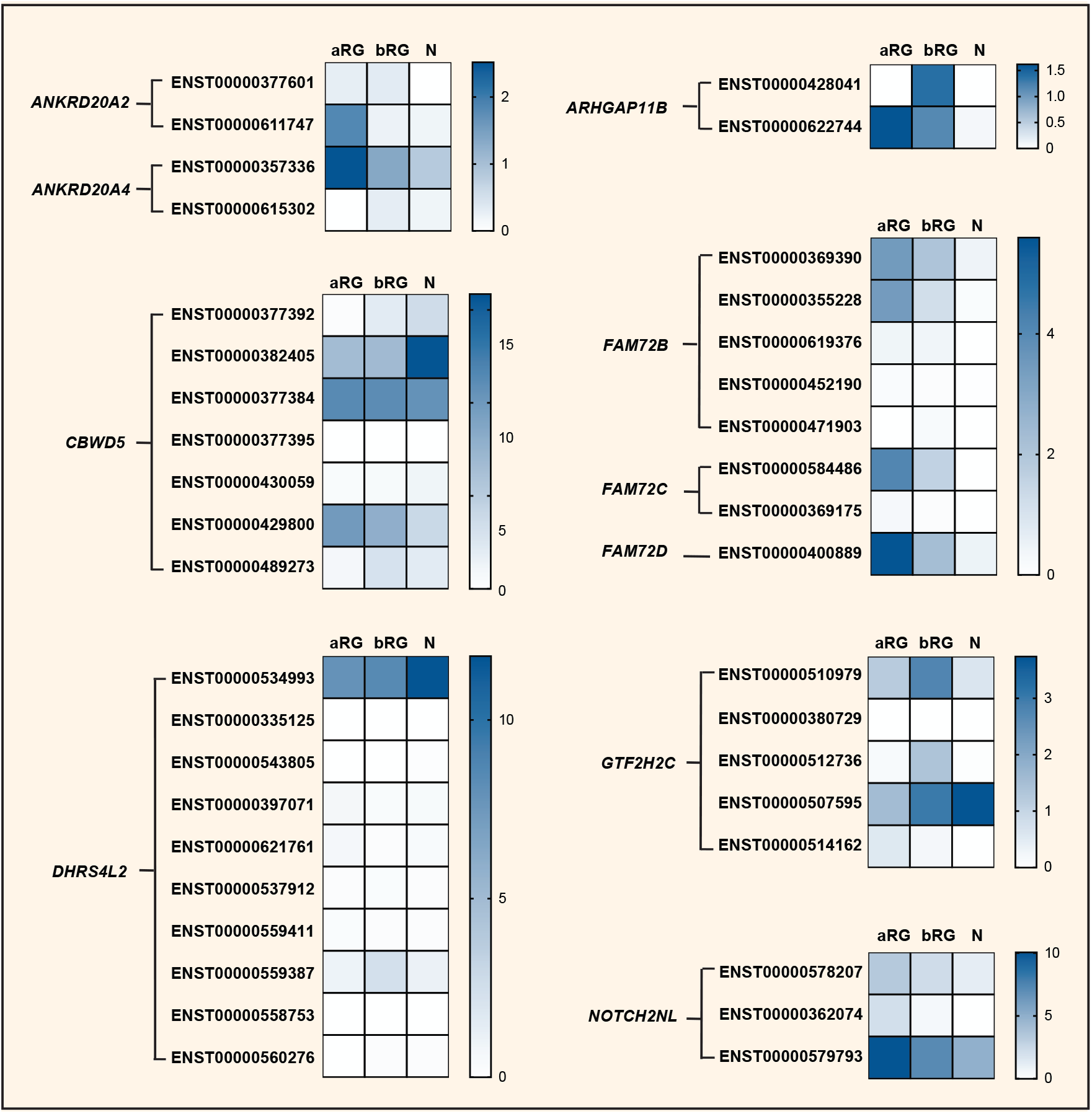
Cell-type specificity of mRNA expression of splice variants encoded by 10 human-specific cNPC-enriched genes. Heatmaps showing TPM expression levels (see color keys) of all protein-coding splice variants encoded by the indicated human-specific cNPC-enriched genes in aRG, bRG and neuron-enriched (N) cell populations from fetal human neocortex (Florio et al., 2015). See Table S4 for mRNA expression data for each cell type and splice variant, including non-coding transcripts. Human-specific genes are grouped based on orthology, and splice variants (indicated by NCBI transcript IDs) encoded by the respective cNPC-enriched human-specific gene(s) are grouped together. Note the specific expression of ENST00000428041, a splice variant of *ARHGAP11B* uniquely expressed in bRG. Splice variant-specific mRNA expression was assessed using the Kallisto algorithm.

## Tables

**Table 1 Primate-specific genes**

**Table S1 cNPC-enriched genes**

**Table S2 qPCR primer**

**Table S3 Primer for ISH probes**

**Table S4 mRNA expression data of splice variants**

## Supplementary files

**File S1 ANKRD20A alignment**

**File S2 ARHGAP11 alignment**

**File S3 CBWD alignment**

**File S4 DHRS4 alignment**

**File S5 FAM72 alignment**

**File S6 GTF2H2 alignment**

**File S7 NOTCH2 alignment**

**File S8 ZNF98 alignment**

